# Reciprocal Best Structure Hits: Using AlphaFold models to discover distant homologues

**DOI:** 10.1101/2022.07.04.498216

**Authors:** Vivian Monzon, Typhaine Paysan-Lafosse, Valerie Wood, Alex Bateman

## Abstract

The conventional methods to detect homologous protein pairs use the comparison of protein sequences. But the sequences of two homologous proteins may diverge significantly and consequently may be undetectable by standard approaches. The release of the AlphaFold 2.0 software enables the prediction of highly accurate protein structures and opens many opportunities to advance our understanding of protein functions, including the detection of homologous protein structure pairs. In this proof-of-concept work, we search for the closest homologous protein pairs using the structure models of five model organisms from the AlphaFold database. We compare the results with homologous protein pairs detected by their sequence similarity and show that the structural matching approach finds a similar set of results. Additionally, we detect potential novel homologues solely with the structural matching approach, which can help to understand the function of uncharacterised proteins and make previously overlooked connections between well-characterised proteins. We also observe limitations of our implementation of the structure based approach, particularly when handling highly disordered proteins or short protein structures. Our work shows that high accuracy protein structure models can be used to discover homologous protein pairs, and we expose areas for improvement of this structural matching approach.

## 2 Introduction

For the last decades the field of similarity searching between proteins has been dominated by sequence similarity methods. This is due to the vast numbers of available protein sequences in databases such as UniProt [1]. With the availability of highly accurate structural models for all proteins [2], structural similarity may begin to supplant or enhance sequence search for some applications. In this manuscript, we investigate the use of structural similarity search to identify homologous protein pairs between human and four model organisms. Identification of homologous functioning proteins between a human and a model organism can help researchers connect experimental data across species and identify relevant model organism proteins and genes for the study of human disease. There are numerous examples where the identification of these connections has advanced molecular biology. For example, in yeast, the eukaryotically conserved KEOPS complex is composed of five subunits (Pcc1p, Kae1p, Bud32p, Cgi121p and Gon7p) and functions as a tRNA modifier [3, 4]. The KEOPS Gon7p subunit was assumed to be fungal specific [5]. However, the human C14orf142 protein was later proved to be a distant Gon7p orthologue [6]. In another example, the worm sup-45 (now called affl-2) protein was shown to be an orthologue of the human AF4/FMR2 family proteins, which are known to be involved in translation elongation [7]. A useful approach to identify potential orthologous proteins between species is the use of reciprocal best hits (RBH) [8]. In this approach, to identify pairs of orthologues between two species an all against all sequence comparison of the two protein sets would be performed, often using BLASTP [9]. A pair (A, B) of orthologues is identified when the best scoring hit of protein A from one organism is protein B in the second organism and reciprocally, the best scoring hit for protein B is protein A. In this manuscript we extend this idea using structure comparison to define reciprocal best structure hits (RBSH) in an analogous way with the aim to identify the closest homologous protein pairs (Figure 1). It is well known that sequence similarity is less conserved than structural similarity [10] and hence this structure based approach should in principle enable discovery of hitherto undiscovered homologous relationships. It is important to note here that we are not using the method to detect orthology relationships, which become less certain as levels of divergence increase. However, it is likely that many of the novel relationships we detect may represent novel orthologues, particularly in the case where there is only a single homologue in each species.

**Figure 1:**
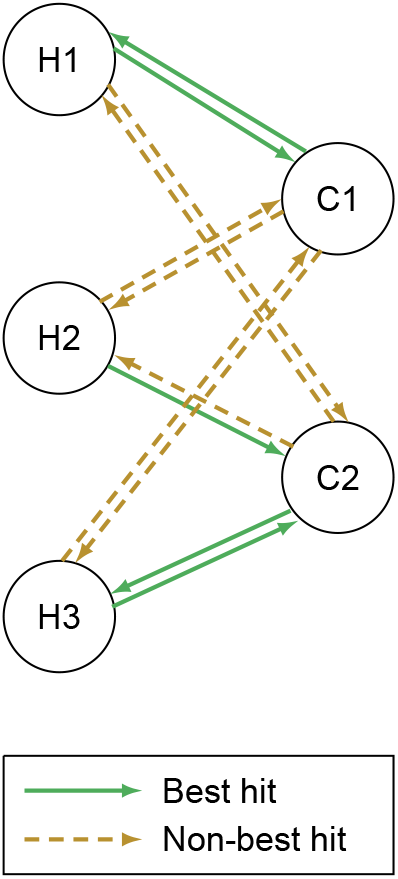
Definition of best reciprocal hits and best reciprocal structure hits: In this scheme protein H1 is the best hit to protein C1 and vice versa. Therefore, H1 and C1 form a reciprocal best hit (RBH) if sequence comparison was used and a best reciprocal structure hit (RBSH) if structure comparison was used. Proteins H3 and C2 form a RBH/RBSH pair while no best reciprocal hit is found for H2. Adapted from [11].

## 3 Methods

We used data from the UniProt Reference Proteomes for the following organisms: *Homo sapiens* (human), *Caenorhabditis elegans* (worm), *Drosophila melanogaster* (fruit fly), *Saccharomyces cerevisiae* (budding yeast) and *Schizosaccharomyces pombe* (fission yeast) [1]. We selected this set of organisms to investigate phylogenetically distant but well-studied species across the opisthokonts.

### 3.1 Protein sequence and structure data sets

We downloaded the AlphaFold structural models for the five selected organisms from the AlphaFold Protein Database (release v2) [12]. Proteins longer than 2,700 residues are only available as multiple fragment entries in the AlphaFold database and were ignored for this study (208 *Homo sapiens* proteins). We extracted the sequences for each of the available structure models in Fasta format. Thus, our data set contains 20,296 protein sequences and structure models for *Homo sapiens* (*H. sapiens*), 19,694 for *Caenorhabditis elegans* (*C. elegans*), 13,458 for *Drosophila melanogaster* (*D. melanogaster*), 6,040 for *Saccharomyces cerevisiae* (*S. cerevisiae*) and 5,128 for *Schizosaccharomyces pombe* (*S. pombe*).

### 3.2 Detection of Reciprocal Best Hits using BLAST

In the Reciprocal Best Hit procedure, an all-against-all BLASTp search is carried out for the two proteomes of interest. Each protein of one proteome is searched against the proteins of the second proteome and vice versa. The BLASTp results in both directions are sorted by E-value and the hit with the lowest E-value is kept as the best hit. In case more than one hit has the lowest E-value, the bit score is used as additional selection criteria, with the highest bit score being selected. The best hits are compared and a reciprocal best hit is identified if the best query target match in one direction matches the best query target match in the other direction.

To detect reciprocal best hits for our study, we carried out a BLAST search (version 2.12.0+) of the *H. sapiens* sequences defined above against each of the *C. elegans, D. melanogaster, S. cerevisiae* and *S. pombe* protein sets and vice versa [9]. We also performed the BLAST search of *S. cerevisiae* sequences against *S. pombe* and vice versa. This pair of yeast species are thought to have diverged 420-330 million years ago, and many orthologous pairs of proteins are highly divergent [13]. The fission yeast proteome appears less rapidly evolving with many proteins more similar to their metazoan orthologs than to the orthologous budding yeast protein [13]. We set an E-value threshold of 0.01 and we required the sequence match to cover 75% of both sequences. Pairs of sequences which passed the E-value and coverage thresholds were sorted by E-value and the hit with the lowest E-value was kept as the best hit, as described above. In cases where the next hit had the same E-value score we kept the hit or hits with the highest bit score [14]. The analysis was performed in both directions, with each organism being both the query proteome and the subject proteome. As described above, if the best hit in one direction was also the best hit in the other direction, the protein pair was kept as a reciprocal best hit (RBH).

### 3.3 Detection of Reciprocal Best Structure Hits using Foldseek

To detect reciprocal best structure hits (RBSH), we carried out structure comparisons with Foldseek (release 1-3c64211 (February 9, 2022)), an extremely fast structure comparison tool [15]. Compared to the other well established structure aligners DALI and TMalign, Foldseek has a lower sensitivity than DALI and a similar accuracy to TMalign. The essential benefit of Fold-seek over DALI and TMalign is its high speed [15]. We compared the human AlphaFold model set against each of the *C. elegans, D. melanogaster, S. cerevisiae* and *S. pombe* models in turn and the *S. cerevisiae* AlphaFold models against the *S. pombe* models using an E-value threshold of 1e-4. To obtain the reciprocal best hits, we used the same methodology as the one used for RBH, the hits by E-value were sorted and the hit with the lowest E-value was kept as best hit. In case the next hit had the same E-value, the hit or hits with the highest bit score was kept. We kept the reciprocal best structure hits, which passed a coverage threshold of 75%.

For a validation of the Foldseek method with regard to highly disordered proteins, we removed protein regions with a confidence pLDDT score below 50 in the AlphaFold structure models. We repeated the detection of RBSH as described above using only the protein regions with a higher confidence prediction score.

### 3.4 Results verification

To verify the results, we used the PANTHER sequence classification files (Release 17.0) for each of the five model organisms [16]. PANTHER provides families of evolutionarily related proteins, usually at the level of the orthologous group. The reciprocal best hits with the same PANTHER family classification were counted as true positive hits and the ones with a different classification or without any classification as false positives. To further verify the results we compared the domain content of the predicted homologues. For the domain comparison, we searched the Pfam HMM-profiles (version 35.0) against the organisms sequences using the HMMER tool (version 3.3.2) with the gathering (GA) threshold option. To identify and assess novel homologies that had not been identified by any other ortholog prediction method we used curated inventories of orthologs between *S. pombe* and *S. cerevisiae, S. pombe* and *H. sapiens*. These ortholog inventories have been manually constructed over a 20 year period from a consensus of multiple ortholog prediction resources, divergent orthologs reported in the literature and directed searches for missing members of conserved complexes [17]. At the beginning of this work ortholog coverage of the fission yeast proteome was already 78.9% for budding yeast and 70.9% for human.

To investigate the effect of the disordered protein fraction on the structural alignment method, we applied IUPred (IUPred2a) on the protein sequences of the used model organisms with the IUPred2-type “long” option for predicting long disordered regions.

## 4 Results

Figure 2 shows the overall number of RBH and RBSH identified between *H. sapiens* and each of the model organisms as well as between *S. cerevisiae* and *S. pombe*. Using Foldseek 4,316 reciprocal best structural hits were found between *H. sapiens* and *D. melanogaster*, 3,837 RBSH between *H. sapiens* and *C. elegans*, 1,921 RBSH between *H. sapiens* and *S. cerevisiae* and 2,095 RBSH between *H. sapiens* and *S. pombe* (Figure 2). These represent 32.07%, 19.69%, 31.80% and 40.85% of the model organisms’ proteome size, respectively. Between the two yeasts *S. cerevisiae* and *S. pombe* 2,751 RBSH were found (Figure 2). More than half of the RBSH found with Foldseek are common with the RBH found with BLASTP (Suppl. table S1).

**Figure 2:**
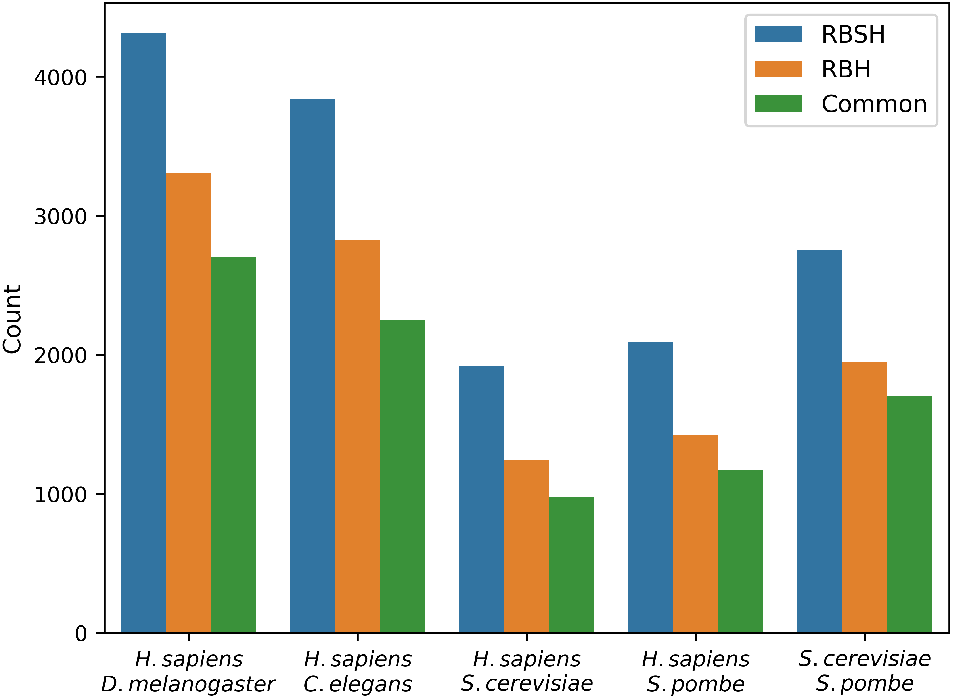
Overview of the detected RBH and RBSH: this bar plot shows the number of RBSH found with Foldseek and RBH found with BLASTP. The common reciprocal hits represent the matches found with both analysis approaches after applying a 75% sequence (RBH) or structure (RBSH) coverage threshold.

The detected RBSH include known homologues such as the human proteasome assembly chaperone 4 (PAC4) (UniProtKB:Q5JS54), which was described as homologous to the POC4 protein in *S. cerevisiae* based on targeted pairwise alignments (UniProtKB: Q12245) [18]. The RBSH approach also found human PAC4 homologues in *S. pombe* (UniProtKB:Q9P7J0), *C. elegans* (UniProtKB:Q9TYS7) and *D. melanogaster* (UniProtKB:Q7JWR4). All of these proteins show a high structural similarity to the *S. cerevisiae* chaperone although sequence similarity was undetected by RBH or any sequence based ortholog predictor.

The RBSH approach also confirms divergent connections made previously based on homology-search algorithms. Wideman connected the human ER-membrane protein complex Emc7 subunit (UniProtKB:Q9NPA0) to the Sop4p protein in *S. cerevisiae* (UniProtKB:P39543) using BLAST and pHMMer [19]. The structural matching approach confirms this finding and provides additional support for the proposed equivalence of these divergent ECM subunits. The RBSH approach also detects the corresponding homologues in *S. pombe* (UniProtKB:O94694), which was previously unconnected to *S. cerevisiae* Sop7p.

We validated the detected RBH and RBSH using the PANTHER classification system [16] and could show that the majority of matches (>90%) share a common PANTHER family classification (Suppl. Figure S1). We calculated the precision of the RBH and RBSH results, by counting the reciprocal matches with the same PANTHER family classification as true positives and the reciprocal matches without the same PANTHER family as false positives. The RBH method yielded higher precision scores compared to the RBSH method (Table 1). We also compared the precision score of the RBSH results before and after applying the coverage threshold of 75% as described in the Method section. The results show that the coverage threshold yields a higher precision (Table 1). Additionally, we tested different coverage thresholds for both the sequence and structural matching approach (Suppl. Figure S2, S3). However, not every protein has a PANTHER classification, with this in mind, it is necessary to investigate whether the RBSH found without the same PANTHER family classification are incorrectly detected by the RBSH method or are novel homologous protein pairs, which still need to be classified in the PANTHER database.

Overall, we see that in each case there are more putative closest homologous pairs found with the structure based method compared to the sequence based method. However, we see that not all of the pairs detected are shared between the RBH and RBSH methods. The fraction of matches in common between the RBH and RBSH methods was not as high as we expected, with many homologues differing between the two approaches. We further analysed the matches missed by each method to understand how much of the difference is based on discrepancies between the two approaches. We observed that many of the matches found as the best hit with the RBSH method were indeed found by the RBH method as a lower scoring match and vice versa. Thus the differences are not due to the ability of each method to find homologous sequences or structures, but more likely due to different pairs of proteins being the best match when viewed through the lense of sequence comparison or structure comparison. We are showing such an example in figure 3, where it seems that the different methods are selecting different members of a post-divergence paralogous pair within an orthologous group. The average number of detected homologous proteins of reciprocal protein pairs is slightly lower for the proteins found as best reciprocal hits with both the RBH and RBSH method (Suppl. table S2). This further strengthens the assumption that the discrepancies are due to the different pairs of best reciprocal hits found by both methods.

**Figure 3:**
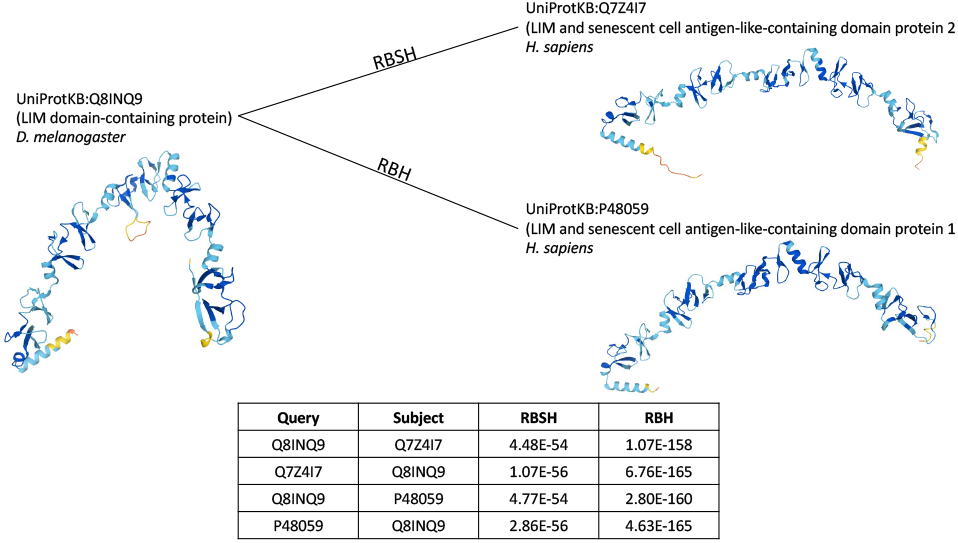
An example of different closest homologues selected with the RBH and RBSH methods: The RBH as well as the RBSH method found more than 20 different homologous proteins for the Drosophila LIM domain-containing protein (UniProtKB:Q8INQ9) in *H. sapiens* with different E-values scores and sequence coverage values. The top ranked protein found with the RBH method, was found with the RBSH method as second best hit and vice versa.

Interesting examples of homologous protein pairs, whose similarity was detected by both approaches, but only found by the RBSH approach as best reciprocal match, include the human transcription factor IIIC subunit GTF3C4 (TFIIIC90) (UniProtKB:Q9UKN8), which was before only based on interaction partners connected to the Sfc9 protein in *S. pombe* (UniProtKB: O13650) [20].

Another new connection is the *S. cerevisiae* nuclear pore localised protein PML39 (UniProtKB:Q03760), a nuclear peripheral protein required for the nuclear retention of unspliced mRNA [21]. PML39 was structurally matched to the *S. pombe* Rsm1 (UniProtKB:O94506). Fission yeast Rsm1 is a poorly characterised C3HC protein with defects in RNA export and a G2/M cell cycle transition (elongated at division) phenotype [22, 23]. The fission yeast Rsm1 is the predicted orthologue of the poorly characterised human ZC3HC1 protein (UniProtKB:Q86WB0). This new connection provides additional support for the reinvestigation of human ZC3HC1 for a potential role in nuclear RNA surveillance or transport.

Both the RBSH and RBH methods found matches exclusive to the method. The number of reciprocal best hits found by one method exclusively, but not detected by the other method, even with lower scoring matches, are listed in Table 2. These RBSH and RBH have a lower sequence similarity, representing potential distantly related homologues (Suppl. Figure S4). A selection of matches exclusively found by the RBSH method are discussed below.

**Table 2:** Matches exclusively found by either the RBSH or RBH method

|  | *H. sapiens:*<br>*D. melanogaster* | *H. sapiens:*<br>*C. elegans* | *H. sapiens:*<br>*S. cerevisiae* | *H. sapiens:*<br>*S. pombe* | *S. cerevisiae:*<br>*S. pombe* |
| --- | --- | --- | --- | --- | --- |
| Only RBSH | 84 | 98 | 143 | 111 | 128 |
| Only RBH | 90 | 54 | 22 | 21 | 40 |

### 4.1 Novel distant homologues

To better understand the performance of the RBSH approach we investigated the best reciprocal hits exclusively identified by the RBSH method. To support that these matches share common characteristics and possible function, we conducted two analyses, (i) identification of their protein families using the PANTHER classification system and (ii) comparison of their annotated protein domains using the Pfam database [24]. The results show that this set of proteins contains many RBSH with the same PANTHER family classification or the same Pfam domain annotation (Figure 4). The fraction of RBSH that are uniquely identified but have no support from PANTHER or Pfam varies between the proteome comparisons. The number of RBSH between *H. sapiens* and the model organisms, which have neither a common Pfam domain annotated nor are classified into the same PANTHER family is 37 for *D. melanogaster*, 35 for *C. elegans*, 24 for *S. cerevisiae* and 23 for *S. pombe* (Figure 4). For *S. cerevisiae* and *S. pombe* it is 24 (Figure 4). It should be noted that having common PANTHER or Pfam annotation is not sufficient to demonstrate closest homology, but it does indicate common ancestry of at least part of the proteins.

**Figure 4:**
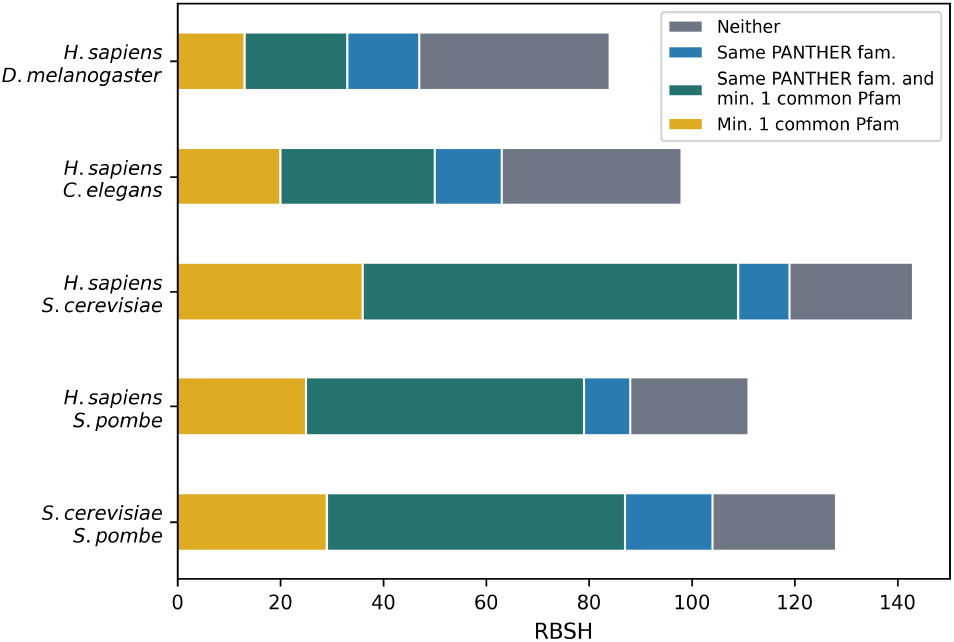
Investigation of the matches exclusively found by the RBSH method: verification of the RBSH exclusively found with Foldseek using the PANTHER classification system and the Pfam database.

We further analysed the RBSH without a common Pfam domain annotation nor PANTHER family classification. These matches would be the potentially most important, representing completely novel homologues identified by the use of structural matching, which can help to understand the function of uncharacterised pro-teins. One example is the human epithelial membrane protein 3 (UniProtKB:P54852), which was found by the RBSH approach to be homologous to an uncharacterised worm protein (UniProtKB:G5EBZ7) (Figure 5a). Another example is the human acid phosphatase type 7 protein (UniProtKB:Q6ZNF0) and an uncharacterised yeast protein (UniProtKB:P53326) (Figure 5b).

**Figure 5:**
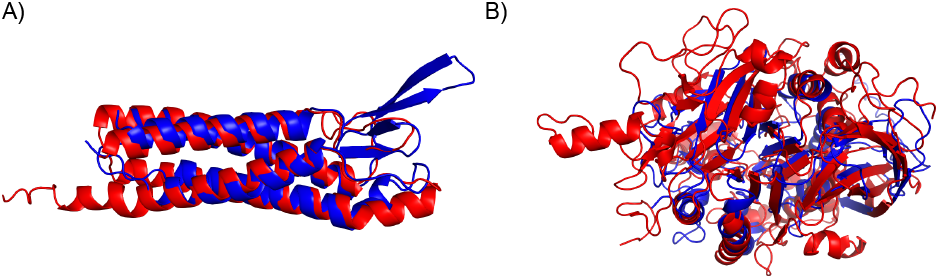
Examples of well superposing RBSH matches without a common Pfam domain annotation nor PANTHER family classification: A) Human epithelial membrane protein 3 (UniProtKB:P54852, in blue) superposed with an uncharacterised worm protein (UniProtKB:G5EBZ7, in red). B) Human acid phosphatase type 7 protein (UniProtKB:Q6ZNF0, in blue) superposed with an uncharacterised yeast protein (UniProtKB:P53326, in red). The structure images were produced using Chimera [27].

We searched for known orthologues for the four described proteins using the ortholog prediction tool DIOPT (version 8.5) [25]. For the *C. elegans* (UniProtKB:G5EBZ7) and *S. cerevisiae* (UniProtKB:P53326) proteins, no orthologous proteins were found in DIOPT. For the *H. sapiens* epithelial membrane protein (UniProtKB:P54852) seven orthologous proteins in *C. elegans* were predicted (UniProtKB:O44789, Q93198, Q9NGJ7, Q11085, Q9N419, Q9NAP4, Q966P3). None of them yielded a better RMSD (Root Mean Square Deviation) value compared to the protein detected with the RBSH method (UniProtKB:G5EBZ7), when superposing the protein structure models using pymol [26]. For the *H. sapiens* acid phosphatase type 7 protein (UniProtKB:Q6ZNF0) two orthologous proteins in *S. cerevisiae* were predicted by DIOPT (UniProtKB:Q12212, Q05924), which indeed yielded a better RMSD score than the protein found by the RBSH method (UniProtKB:P53326), when superposing the structure models.

The following three examples represent potential interesting novel RBSH without a common Pfam domain annotation nor PANTHER family classification in *S. pombe*. First, the structural matching approach detected a relationship between *S. pombe* SPAC323.03c (UniProtKB:Q9UT96) and *S. cerevisiae* PEX8 (UniProtKB:P53248) (Figure 6a,b). No family membership nor detectable sequence similarity outside the fission yeast clade was recorded for *S. pombe* SPAC323.03c using multiple methods [17].

**Figure 6:**
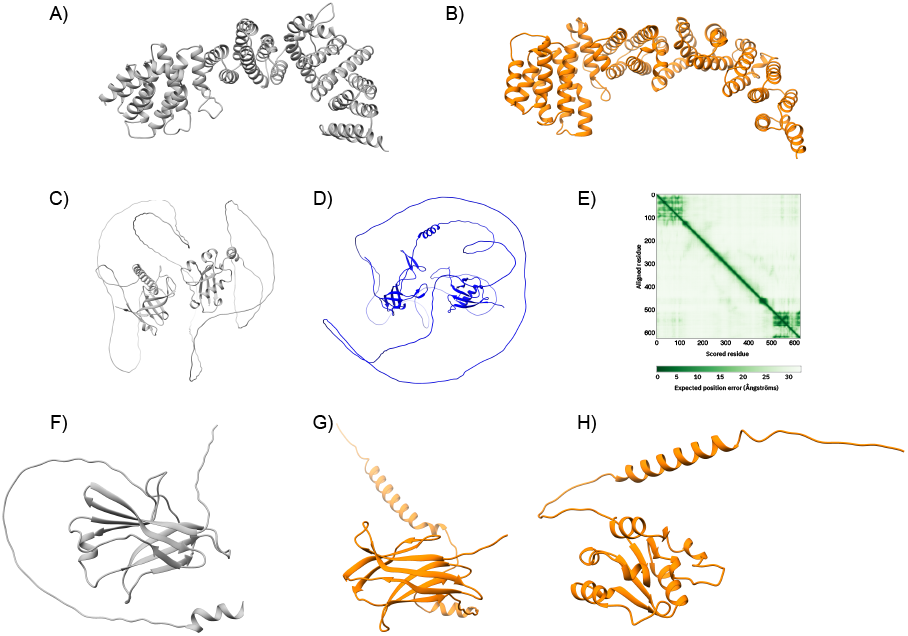
Examples of potential novel homologues in *S. pombe*: The first example links A) *S. pombe* SPAC323.03c (UniProtKB:Q9UT96) with B) *S. cerevisiae* PEX8 (UniProtKB: P53248). The second example predicts C) the *S. pombe* Mug174 protein (UniProtKB:O74434) as structural homologue to D) the human coilin protein (UniProtKB:P38432). E) Predicted aligned error for S. pombe Mug174 AlphaFold structure model. The third example shows a closer structural similarity between F) *S. pombe* SPAC19D5.02c (UniProtKB:Q1K9B6) and G) *S. cerevisiae* EMC10 (UniProtKB:Q12025) rather than with h) *S. cerevisiae* Pex22 (UniProtKB:P39718). The structure images were produced using Chimera [27].

Second, despite over 30 years of intensive study, no fungal orthologue of coilin, a protein important for snRNP biogenesis, has so far been identified [28]. The RBSH approach predicts the uncharacterised *S. pombe* Mug174 (UniProtKB:O74434) as structural homologue of human coilin (UniProtKB:P38432) (Figure 6c,d). Mug174 interacts with four subunits of MTREC, a nucleolar complex functionally connected to snRNA processing [29]. Like Coilin, Mug174 is composed of two distal subdomains, which are not predicted to interact with each other (Figure 6e). The mouse coilin knockout has a strong detrimental effect on fertility, and the fission yeast deletion affects spore number, a common phenotype of meiotic defects [30]. Taken together the interactions, phenotypes and RBSH data demonstrate a compelling functional connection between these two proteins.

The third RBSH is *S. pombe* SPAC19D5.02c (UniProtKB:Q1K9B6) with *S. cerevisiae* EMC10, a subunit of the ER membrane complex involved in the insertion of newly synthesised proteins into the ER membrane (UniProtKB:Q12025) (Figure 6f,g). This finding refutes the previous orthology assignment between *S. pombe* SPAC19D5.02c and *S. cerevisiae* Pex22 (UniProtKB:P39718) (Figure 6h) [17]. The RBSH match is supported by the PANTHER family PTHR21397 which includes the human EMC10 and the *S. pombe* SPAC19D5.02c. Jackhammer using human EMC10 retrieves both *S. cerevisiae* EMC10 and fission yeast SPAC19D5.02c. This finding increases the EMC complex complement in *S. pombe* to eight.

### 4.2 Limitations of the RBSH approach

Within the set of RBSH without a common Pfam domain or PANTHER family classification we observed RBSH that are unlikely to be homologous and were most likely found by the RBSH method due to a high fraction of protein being disordered. Indeed, the percentage of exclusive RBSH matches, but also exclusive RBH matches, slightly increases with a higher disordered protein fraction (Suppl. figure S5). One example is the human endonuclease subunit SLX4 (UniProtKB:Q8IY92) found by the RBSH approach to be homologous to an uncharacterised protein from the fruit fly (UniProtKB:Q9VHT6) (Figure 7). The human endonuclease subunit SLX4 was detected as best hit for the fruit fly protein with an E-value of 9.714E-08 and a bit score of 374 and vice versa the fruit fly protein was selected as best hit with an E-value of 5.890E-10 and a bit score of 455.

**Figure 7:**
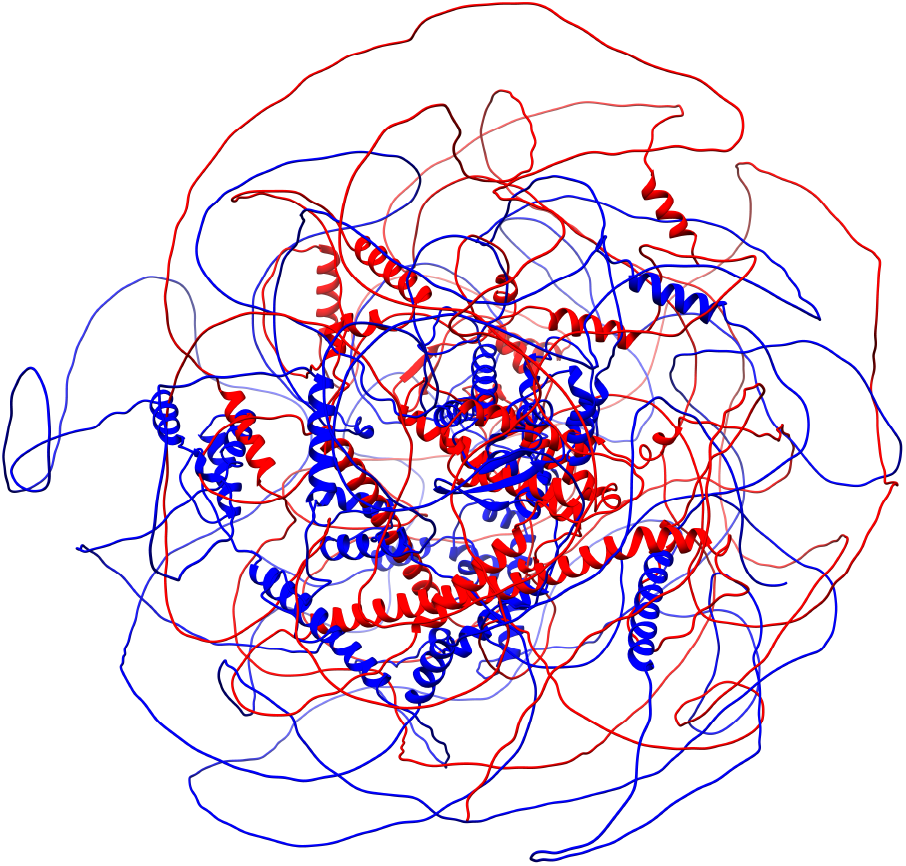
Example of a RBSH unlikely to be homologous: superposition of the human endonuclease subunit SLX4 (UniProtKB:Q8IY92, in blue) and the uncharacterised protein from the fruit fly (UniProtKB:Q9VHT6, in red). The structure image was produced using Chimera [27].

The disordered protein regions largely overlap with low confidence scoring regions [2, 31]. To avoid the detection of RBSH based on disordered regions, we removed the low confidence scoring regions in the AlphaFold structure models. We repeated the detection of RBSH as well as the validation of the results by calculating the precision scores based on the PANTHER classification. The described RBSH shown in figure 7, for example, is no longer detected as a RBSH, suggesting that removing low confidence regions could reduce false positives. But overall, the precision scores dropped or remained the same when comparing it to the RBSH approach without a confidence score threshold (Table S3, Figure S6).

In addition to RBSH exclusively found with the structural matching approach, there are also RBH exclusively found by sequence similarity (Table 2). These matches may represent important failure modes from the RBSH method. We studied the RBH exclusive hits and deduced that potential reasons for the structural matching approach to miss these RBH could be differences in the inter-domain structure orientations, a lower AlphaFold model quality or again a high disordered protein fraction (Figure 8a,b).

**Figure 8:**
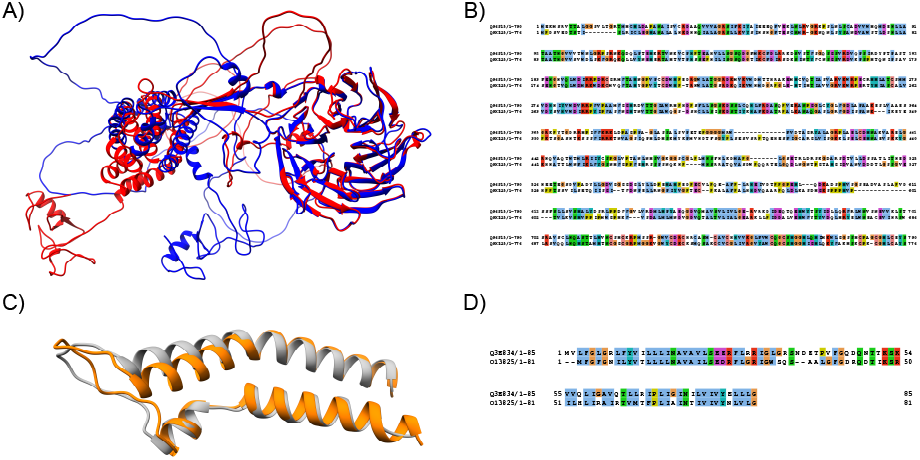
Example of RBH exclusively found by BLASTP: A) Structure model superposition and B) sequence alignment of the human GATOR complex protein WDR24 (UniProtKB:Q96S15) with the fruit fly GATOR complex protein WDR24 (UniProtKB:Q9XZ25). C) Structure model superposition and D) sequence alignment of the transport protein YOS1 of *S. cerevisiae* (UniProtKB:Q3E834) and *S. pombe* (UniProtKB:O13825). The structure images were produced using Chimera and the sequence alignment was performed using Jalview [27, 32]

Lastly, we observed that very short well superposed protein structures are missed, potentially because these protein structures do not achieve the predefined significance threshold due to their small protein size (Figure 8c,d). Indeed, many of the RBH exclusively found by BLASTP have an average length below 200 residues (Suppl. figure S7).

## 5 Discussion

Protein sequences encode the information about protein structures and functions, and the use of protein sequences for the detection of closest homologous protein pairs is widely established [14, 33]. The release of the AlphaFold 2.0 structure predictor gives us now the opportunity to use highly accurate protein structure predictions to find the closest homologous protein pairs [34, 12]. A structural matching approach is expected to detect novel homologues, which were previously missed due to sequence divergence.

In this study, we assess the detection of homologous protein pairs using protein structures in comparison to protein sequences. We used the structure aligner Foldseek to detect RBSH and the sequence aligner BLASTP to detect RBH [15, 9]. Compared to using all best hits, selecting the reciprocal best hits represents a higher barrier for finding homologous proteins and yields a higher precision (Suppl. table S4). A large number of commonly detected protein pairs demonstrates the ability of the structural matching approach. Additionally to commonly detected protein pairs, we found many cases of different protein pairs scored as best reciprocal hits by one approach, which were nevertheless also found by the other approach within the top ranked hits. The comparison between the two approaches is therefore limited by the scoring discrepancies. Lastly, we also found protein pairs exclusively detected by the RBSH or RBH approach.

We used the PANTHER classification system to validate the RBSH and RBH approaches. Even though we found a higher number of RBSH compared to RBH, the validation showed that the conventional RBH approach yields a higher precision score compared to the RBSH approach. This outcome can be explained by limitations of the RBSH approach, but also possibly due to the bias that homologues were so far mainly detected using the protein sequences and protein pairs with similar sequences are therefore better studied and more prevalent in the PANTHER database. It is likely that the PANTHER database aligns well with RBH because it is also a sequence based method [16].

We encountered different limitations of the RBSH approach while investigating the protein pairs exclusively detected by the RBH approach. The structure of highly disordered proteins turned out to be very challenging to align as the RBSH approach on one hand aligned protein structures solely based on the disordered regions, but on the other hand missed homologous proteins due to their disordered regions. When removing the structure regions with a low AlphaFold confidence score, the results improved with regard to single false positive RBSH, which were no longer detected, but overall, the precision score worsened. A possible explanation is that the AlphaFold confidence scores are not homologous between the potential RBSH and consequently removing different parts of the two proteins forming a true positive RBSH pair, can lead to a less significant E-value and lower structure coverage. Furthermore, homologous protein pairs were missed by the RBSH approach due to different domain structure orientations or due to a small protein size, so that the structural matches are not able to achieve statistical significance.

When investigating the protein pairs detected with the RBSH approach, we detected potential novel homologues, which can be particularly useful for describing uncharacterised proteins and to draw new functional connections between different species. Model organisms are generally well studied and hence the majority of proteins of their proteomes are already characterised to some degree [35]. Nevertheless, we showed that the structural matching approach can find potential novel homologues, confirm previously made predictions or even correct mistaken orthologies. Importantly these new connections can provide functional clues for previously unknown or partially characterised proteins. Even for *S. pombe* a species with extensive coverage of curated orthologues the RBSH approach provided 39 novel *S. cerevisiae* and 11 novel human homologues, which were added to the PomBase database (Suppl. Tables S5, S6) [36]. Most of these novel connections provided additional functional information, or supported existing knowledge for poorly characterised proteins. We believe that the RBSH approach can advance the understanding of protein functions across the model organisms and also for many other species, as for example, emerging pathogens [37].

## 6 Conclusion

We have introduced a novel extension of the reciprocal best matching method to use structural models. Our work shows that such a method works in principle and has a similar set of results to sequences based on RBH detection. Interestingly the structure based search when faced with many choices in large paralogous families often selects a different pair of best hits compared to the sequence based approach. We also find that there are a small set of unique matches of potential distant homologues that can be discovered by the RBSH method. We do find some deficiencies in our implementation of the RBSH method. For example, the method seems to be confused by highly disordered proteins and also often fails to find obvious homologues for short proteins.

We hope that our work will encourage others to investigate whether structural models can be used for tasks that have traditionally been the preserve of sequence based methods. We also see many directions for the refinement of the RBSH method through the use of alternative structural comparison methods and filtering procedures. For example, flexible structure matching may enable RBSH identification even when interdomain orientations have been incorrectly predicted. We also envisage combined methods that make use of both sequence and structural information to infer homology across longer evolutionary distances.

## Supporting information

Supplementary tables and images will be used for the link to the file on the preprint site.

## 7 Author contributions statement

V.M. performed the search for RBH and the verification of detected RBH and RBSH. T.P-L. conducted the search for RBSH and contributed to the assessment of detected RBSH. V.W. conducted the extensive assessment of detected RBSH and identification of novel homologues between *S. pombe* and *S. cerevisiae* and between *S. pombe* and *H. sapiens*. A.B. conceived and supervised the study and contributed to the assessment of detected RBSH. T.P-L. and V.W. contributed to the writing of this manuscript. V.M. and A.B. were major contributor in writing this manuscript. All authors read and approved the final manuscript.

## 8 Data availability

The results data of the detected RBSH and RBH can be found in an institutional repository of the University of Cambridge [38]. This repository also includes the list of RBSH, which were added to PomBase, with additional description. The code to find RBSH and RBH is provided in a GitHub repository https://github.com/VivianMonzon/Reciprocal_Best_Structure_Hits.

## 9 Acknowledgments

V.M., T.P-L. and A.B. were supported by core EMBL funding. VW was supported by Wellcome Trust [Grant number 218236/Z/19/Z].

