## Supplementary tables and images will be used for the link to the file on the preprint site. for "Reciprocal Best Structure Hits: Using AlphaFold models to discover distant homologues"

**Table S1: Results overview:** Number of detected RBSH, RBH and commonly found closest homologous protein pairs between *H. sapiens* and the four model organisms as well as between *S. cerevisiae* and *S. pombe*.

| | *H. sapiens* - *D. melanogaster* | *H. sapiens* - *C. elegans* | *H. sapiens* - *S. cerevisiae* | *H. sapiens* - *S. pombe* | *S. cerevisiae* - *S. pombe* |
|---|---|---|---|---|---|
| RBSH | 4,316 | 3,837 | 1,921 | 2,095 | 2,751 |
| RBH | 3,306 | 2,824 | 1,245 | 1,423 | 1,949 |
| Common | 2,706 | 2,251 | 979 | 1,175 | 1,706 |

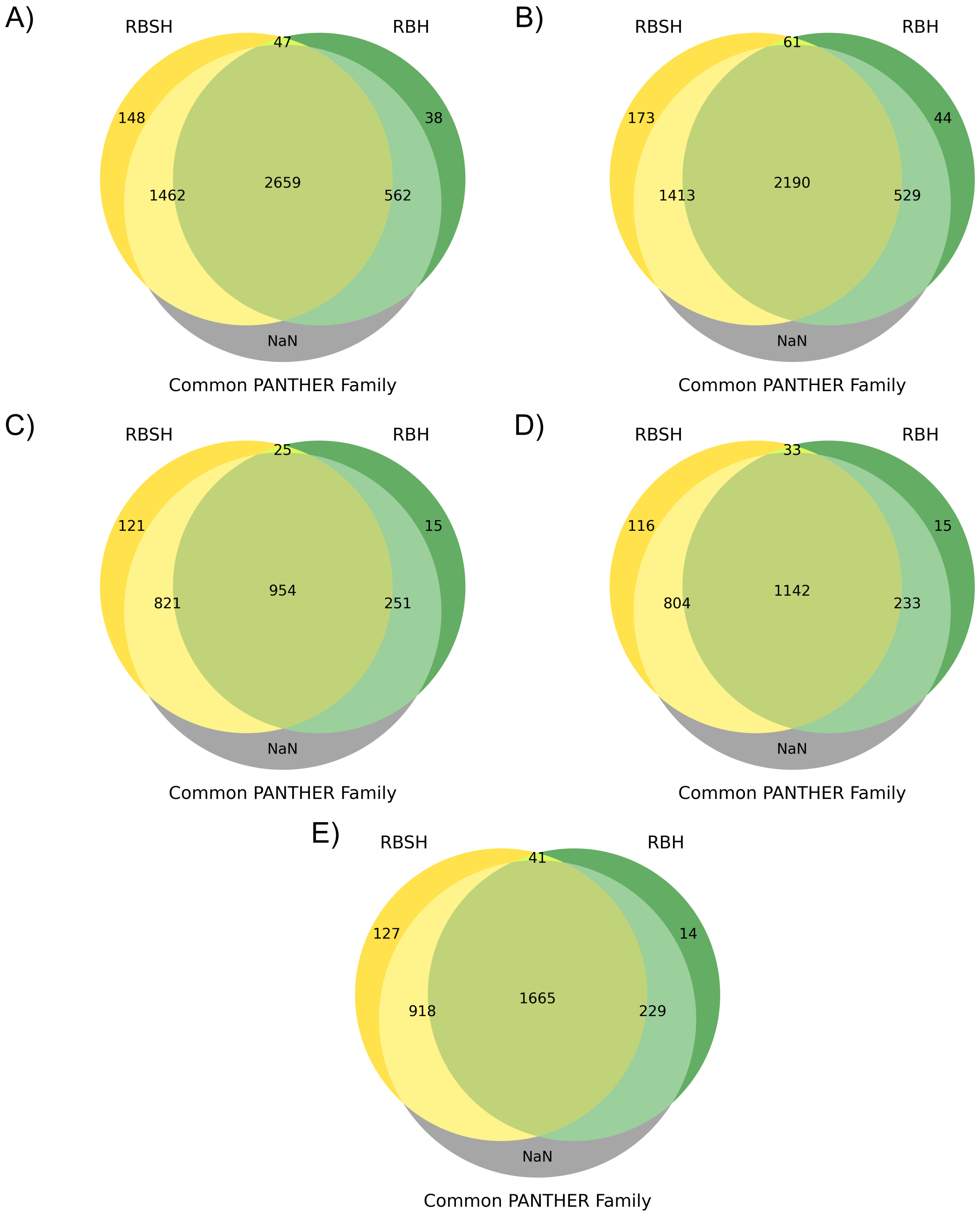


**Figure S1: Comparison of RBH and RBSH methods with PANTHER family classification:** comparison between RBSH detected with Foldseek and RBH detected with BLASTP for A) *H. sapiens* and *D. melanogaster* B) *H. sapiens* and *C. elegans*, C) *H. sapiens* and *S. cerevisiae*, D) *H. sapiens* and *S. pombe*, and E) *S. cerevisiae* and *S. pombe*. 'Common Panther Family' represents how many of the RBH or RBSH found with the two approaches have a common PANTHER family classification. The number of protein pairs with a common PANTHER Family classification were not counted ('NaN').

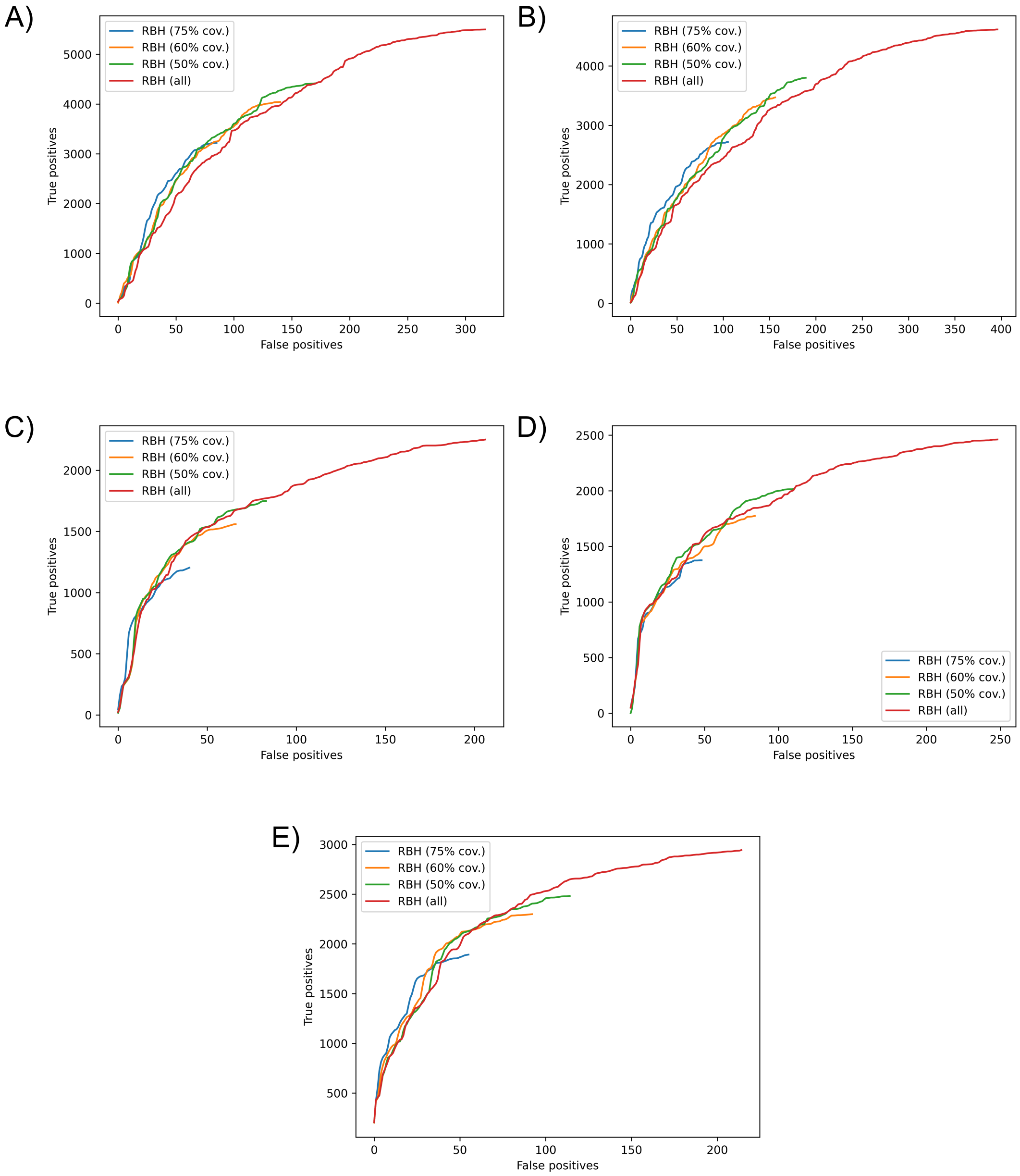


**Figure S2: Investigating different coverage thresholds for the RBH approach:** Validation of the RBH method testing different coverage thresholds. 'True positives' represents the protein pairs with the same PANTHER family, and 'False positives' represents the protein pairs without the same PANTHER family or without any PANTHER classification. The results are shown for A) *H. sapiens* and *D. melanogaster*, B) *H. sapiens* and *C. elegans*, C) *H. sapiens* and *S. cerevisiae* D) *H. sapiens* and *S. pombe* and E) *S. cerevisiae* and *S. pombe*.

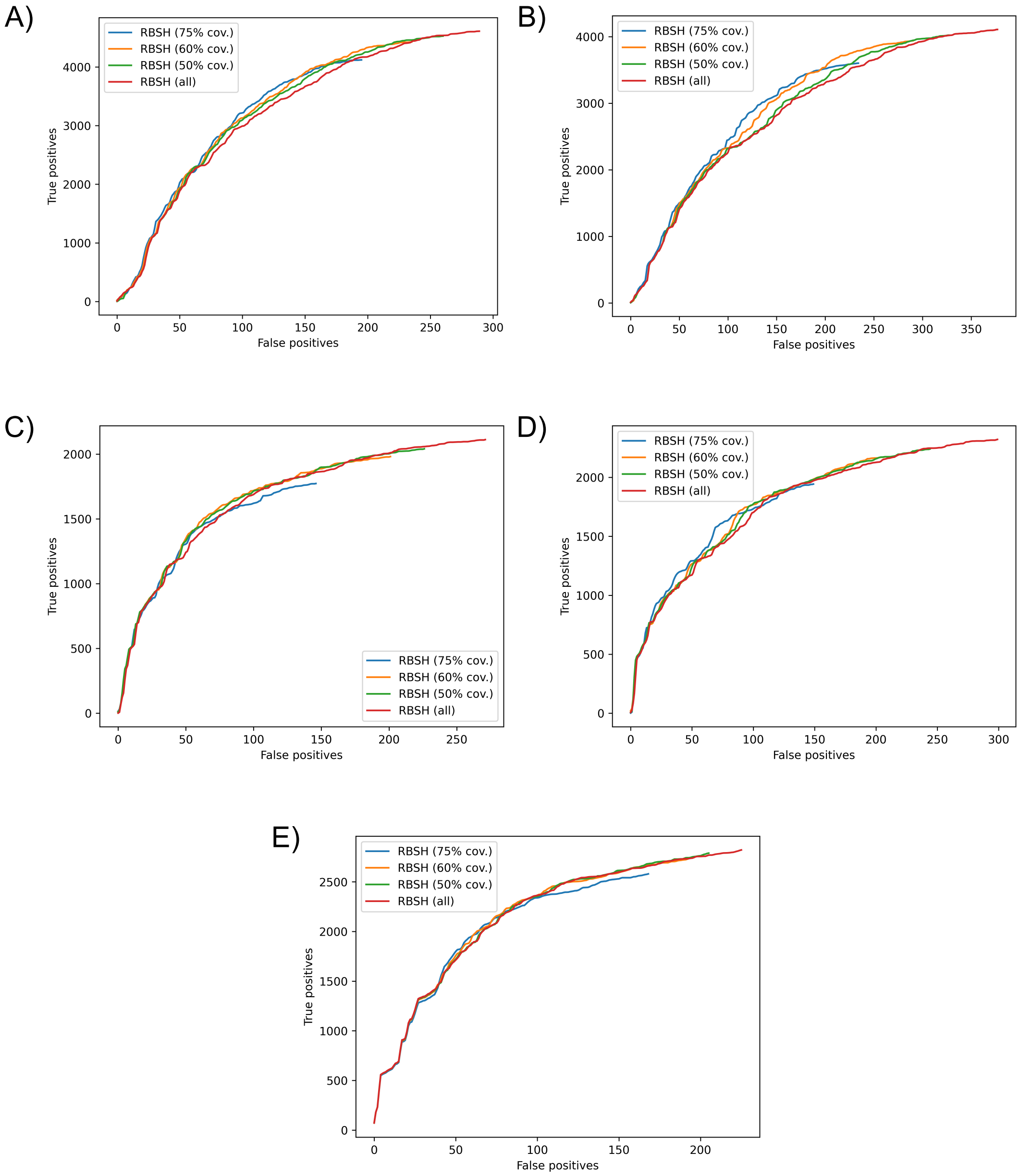


**Figure S3: Investigating different coverage thresholds for the RBSH approach:** Validation of the structural matching approach testing different coverage thresholds. 'True positives' represents the protein pairs with the same PANTHER family, and 'False positives' represents the protein pairs without the same PANTHER family or without any PANTHER classification. The results are shown for A) *H. sapiens* and *D. melanogaster*, B) *H. sapiens* and *C. elegans*, C) *H. sapiens* and *S. cerevisiae* D) *H. sapiens* and *S. pombe* and E) *S. cerevisiae* and *S. pombe*.

**Table S2: Number of homologous proteins found on average:** the table lists the number of homologous proteins found on average per query protein representing the different number of choices on average to form a reciprocal best hit (RBH) or reciprocal best structure hit (RBSH). The results distinguish all proteins found in a reciprocal hit (all) and proteins found by both the RBH and RBSH methods (common).

| | *H. sapiens* (p1) - *D. melanogaster* (p2) | | *H. sapiens* (p1) - *C. elegans* (p2) | | *H. sapiens* (p1) - *S. cerevisiae* (p2) | | *H. sapiens* (p1) - *S. pombe* (p2) | | *S. cerevisiae* (p1) - *S. pombe* (p2) | |
|---|---|---|---|---|---|---|---|---|---|---|
| | p1 | p2 | p1 | p2 | p1 | p2 | p1 | p2 | p1 | p2 |
| RBH all | 4.77 | 5.97 | 4.44 | 5.34 | 2.77 | 3.99 | 2.52 | 3.83 | 2.09 | 2.23 |
| RBH common | 2.42 | 2.82 | 2.53 | 2.99 | 1.99 | 2.83 | 2.03 | 2.93 | 1.78 | 1.85 |
| RBSH all | 9.01 | 12.72 | 18.16 | 20.4 | 6.26 | 10.72 | 6.0 | 10.85 | 5.94 | 6.36 |
| RBSH common | 5.28 | 6.71 | 5.9 | 6.89 | 3.51 | 5.88 | 3.93 | 6.83 | 4.9 | 5.11 |

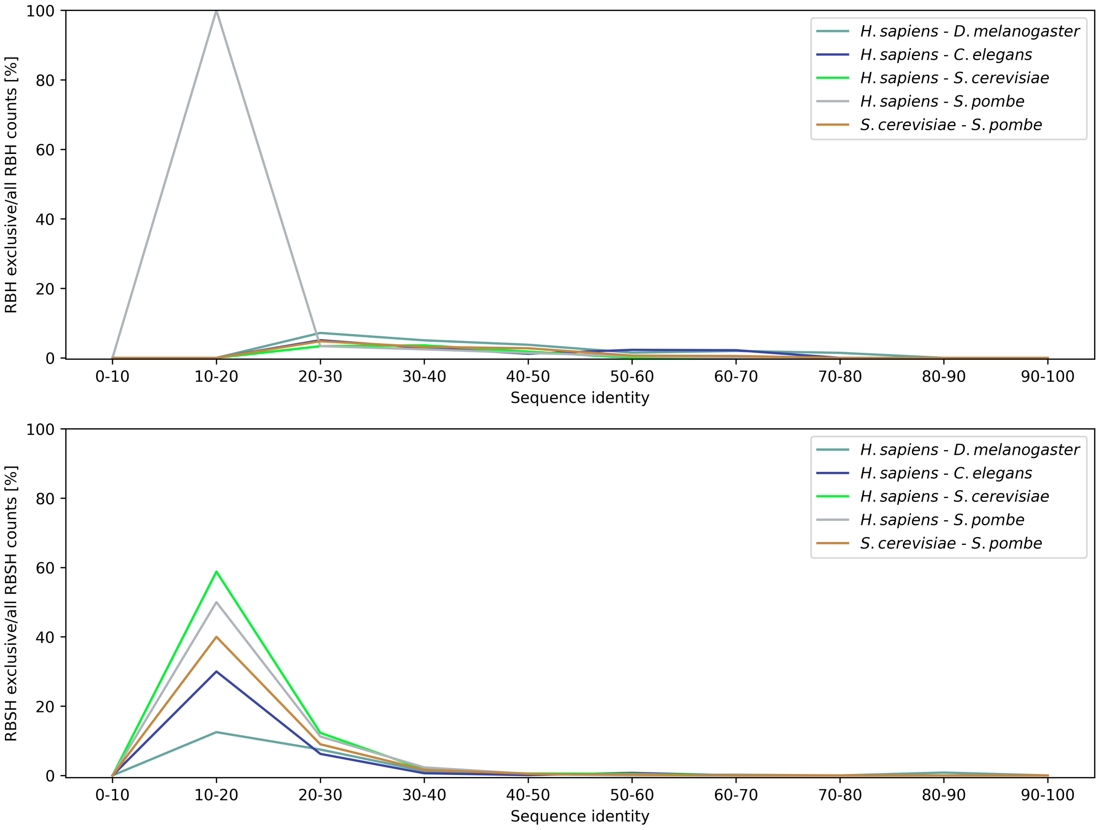


**Figure S4: Sequence similarity of protein pairs exclusively found by the RBH or RBSH method:** Number of protein matches found exclusively by one approach normalised by the total number of matches found by this approach, plotted against the sequence identity calculated between the protein pairs. Analysis results for the different model organisms are differentiated between the RBH approach (plot above) and RBSH approach (plot below).

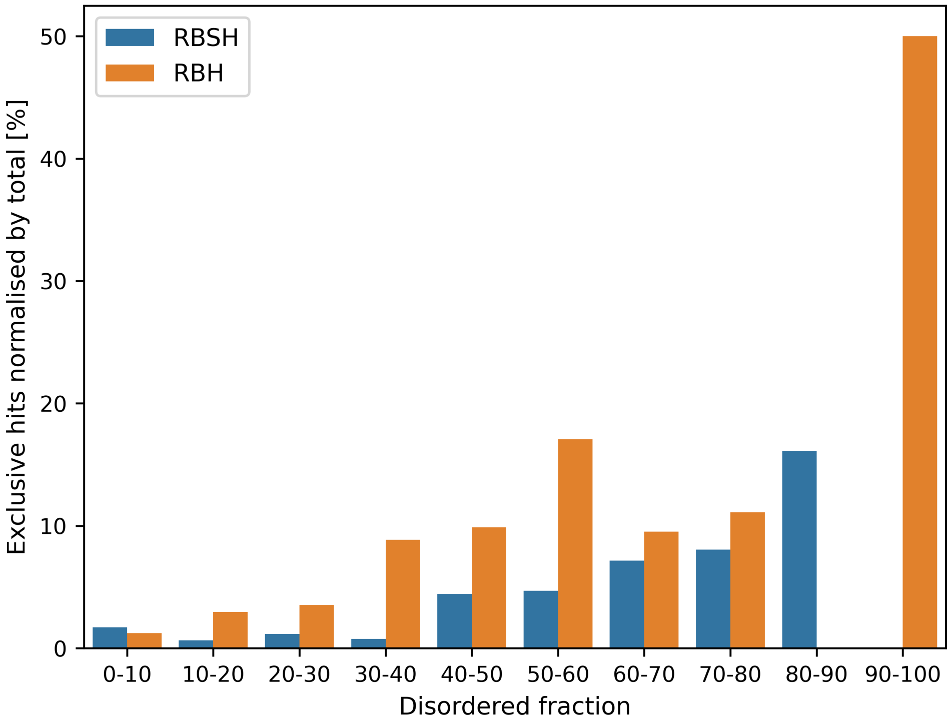


**Figure S5: Percentage of protein matches exclusively found by the RBSH or RBH method per disordered protein fraction:** This plot shows the results of the protein matches between *H. sapiens* and *D. melanogaster*, exemplary for the results with the other model organisms.

**Table S3: Precision score comparison:** Comparison of RBSH approach results with original structure models and only high confidence regions of the structure models (>50 pLDDT score). Precision scores calculated based on PANTHER classification.

| | *H. sapiens* - *D. melanogaster* | *H. sapiens* - *C. elegans* | *H. sapiens* - *S. cerevisiae* | *H. sapiens* - *S. pombe* | *S. cerevisiae* - *S. pombe* |
|---|---|---|---|---|---|
| RBH | 0.974 | 0.969 | 0.973 | 0.968 | 0.973 |
| RBSH (75% cov.) | 0.955 | 0.939 | 0.924 | 0.929 | 0.939 |
| RBSH (no cov. thres.) | 0.941 | 0.916 | 0.886 | 0.886 | 0.926 |
| RBSH confidence score >50 (75% cov.) | 0.955 | 0.932 | 0.911 | 0.917 | 0.938 |
| RBSH confidence score >50 (No cov. thres.) | 0.942 | 0.91 | 0.873 | 0.881 | 0.926 |

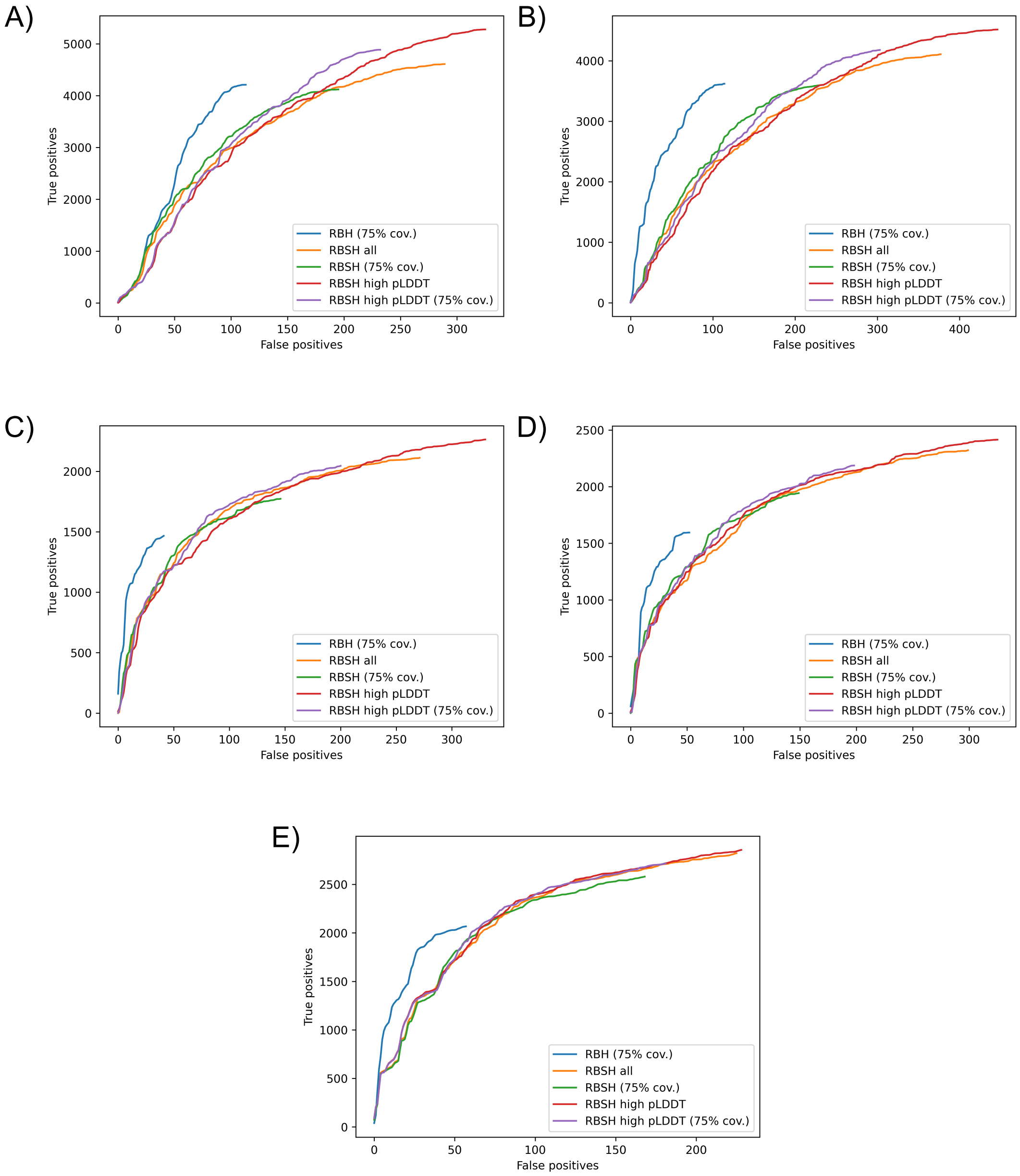


**Figure S6: Validation plots with results of the RBSH approach using pdb files without low confidence regions:** validation plots as shown in figure S2 plus the validation results of RBSH methods using the adapted pdb files. These pdb files do not compromise regions with an AlphaFold confidence prediction below 50 (pLDDT score). 'True positives' represents the protein pairs with the same PANTHER family, and 'False positives' represents the protein pairs without the same PANTHER family or without any PANTHER classification. The results are shown for A) *H. sapiens* and *D. melanogaster*, B) *H. sapiens* and *C. elegans* C) *H. sapiens* and *S. cerevisiae* D) *H. sapiens* and *S. pombe* and E) *S. cerevisiae* and *S. pombe*.

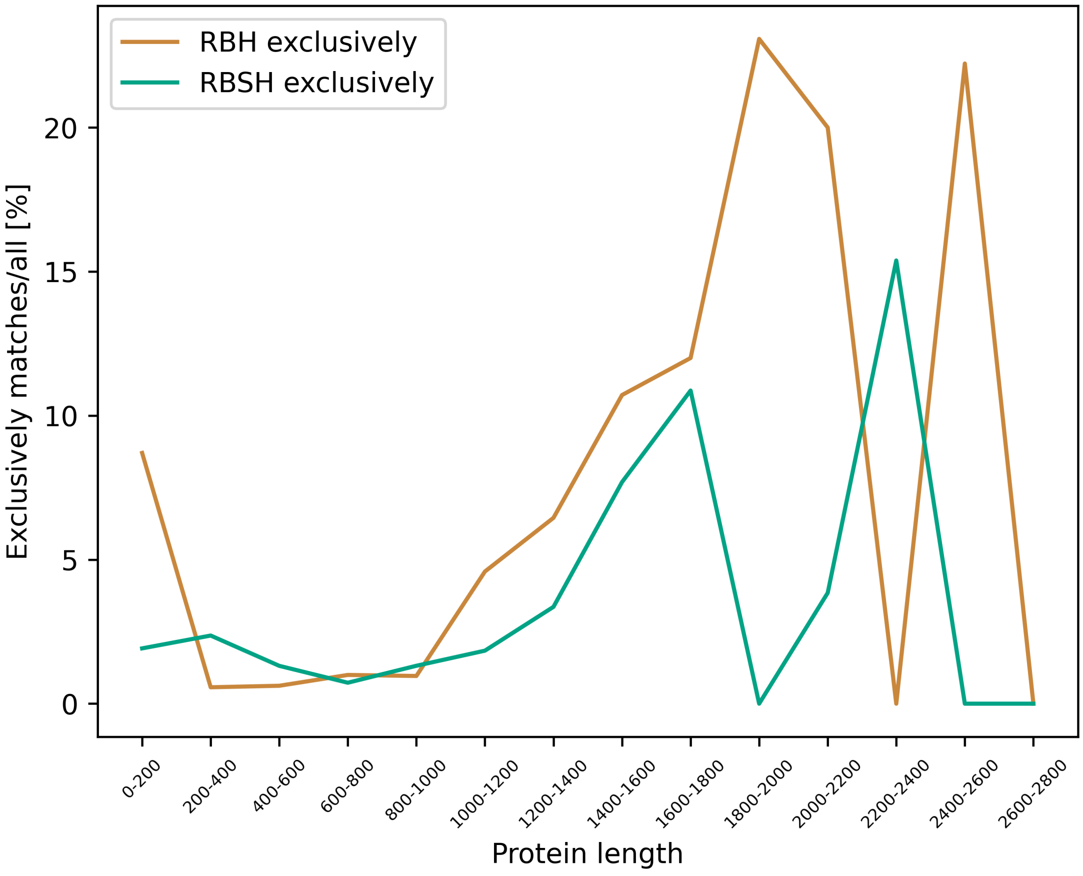


**Figure S7: Percentage of protein matches exclusively found by the RBSH or RBH method per protein length:** this plot shows the results for the protein matches between *H. sapiens* and *D. melanogaster*, exemplary for the results with the other model organisms.

**Table S4: Comparison between the best structure hit and reciprocal best structure hit**: The precision score between the RBSH and best structure hits are compared for the structural matching approach. Precision scores are calculated based on the PANTHER classification. For the best structure hits, the precision is calculated for the best hits selected in both directions, meaning both organisms being once the query proteome, and the average score is listed here.

| | RBSH | Best structure hit (average) |
|---|---|---|
| *H. sapiens*: *D. melanogaster* | 0.955 | 0.707 |
| *H. sapiens*: *C. elegans* | 0.939 | 0.613 |
| *H. sapiens*: *S. cerevisiae* | 0.924 | 0.583 |
| *H. sapiens*: *S. pombe* | 0.929 | 0.605 |
| *S. cerevisiae*: *S. pombe* | 0.939 | 0.844 |

**Table S5: AlphaFold RBSH predictions added to PomBase curated orthologs:** Detected between *S. pombe* and *H. sapiens*.

| *S. pombe* | | | *H. sapiens* | |
|---|---|---|---|---|
| Systematic identifier | Gene symbol | UniProt | HGNC appoved name | UniProt |
| SPAC12B10.02c | pho86 | Q10436 | NAT8 | Q9UHE5 |
| SPCC1682.03c | mug174 | O74434 | COIL | P38432 |
| SPAP27G11.02 | mgr3 | Q9P7N6 | TTC19 | Q6DKK2 |
| SPCC1753.02c | git3 | O94744 | GPR101 | Q96P66 |
| SPAC1952.09c | ach1 | Q9UUJ9 | OXCT1 | P55809 |
| SPCC1393.14 | ten1 | P0C5Y7 | TEN1 | Q86WV5 |
| SPAC1952.08c | | Q9UUK0 | CREG1 | O75629 |
| SPAC2F3.01 | imt1 | O14084 | A4GALT | Q9NPC4 |
| SPAC922.05c | | Q9URX1 | UNC93A | Q86WB7 |
| SPBC21.02 | rtc5 | O94644 | MEAK7 | Q6P9B6 |
| SPBC216.03 | | Q9Y7K0 | BLVRB | P30043 |

**Table S6: AlphaFold RBSH predictions added to PomBase curated orthologs:** Detected between *S. sapiens* and *S. cerevisiae*

| *S. pombe* | | | *S. cerevisiae* | |
|---|---|---|---|---|
| Systematic identifier | Gene symbol | UniProt | SGD gene name & locus ID | UniProt |
| SPAC1B3.10c | | O13875 | HRD3/YLR207W | Q05787 |
| SPBC29A3.17 | gef3 | O59679 | FUS2/YMR232W | Q05670 |
| SPAC13G6.08 | fzr2 | Q09786 | AMA1/YGR225W | P50082 |
| SPCC306.05c | ins1 | Q9Y7R5 | NSG1/YHR133C | P38837 |
| SPBC19G7.08c | art1 | O42956 | RGL1/YPL066W | Q12194 |
| SPBC1289.06c | ppr8 | O94615 | RMD9/YGL107C | P53140 |
| SPBC16G5.16 | | O60130 | AGS1/YIL130W | P40467 |
| SPCP25A2.03 | tho1 | Q9URT2 | HPR1/YDR138W | P17629 |
| SPAC3G9.17 | new8 | G2TRL9 | PPT2/YPL148C | Q12036 |
| SPCC126.08c | | O94401 | UIP5/YKR044W | P36137 |
| SPAC1002.07c | | P79081 | HPA2/YPR193C | Q06592 |
| SPCC1672.10 | mis16 | O94244 | HAT2/YEL056W | P39984 |

| | | | | |
|---|---|---|---|---|
| SPAC1952.12c | csn71 | Q9UUJ7 | CSN9/YDR179C | Q03981 |
| SPAC3C7.04 | | O14130 | PUT3/YKL015W | P25502 |
| SPAC8E11.04c | | O42881 | YLR118C | Q12354 |
| SPBP19A11.06 | lid2 | Q9HDV4 | ECM5/YMR176W | Q03214 |
| SPAC1399.05c | toe1 | Q9HE16 | URC2/YDR520C | Q04411 |
| SPBC19C2.04c | ubp11 | Q9UUD6 | UBP16/YPL072W | Q02863 |
| SPAC323.03c | | Q9UT96 | PEX8/YGR077C | P53248 |
| SPCC1753.05 | rsm1 | O94506 | PML39/YML107C | Q03760 |
| SPAC12G12.12 | gms2 | Q09875 | YPL264C | Q08980 |
| SPAC17C9.16c | mfs1 | Q10487 | FLR1/YBR008C | P38124 |
| SPAC23E2.01 | fep1 | Q10134 | GZF3/YJL110C | P42944 |
| SPBC16E9.19 | pac3 | Q9C0X2 | IRC25/YLR021W | Q07951 |
| SPCC24B10.16c | pac4 | Q9P7J0 | POC4/YPL144W | Q12245 |
| SPBC8D2.07c | sfc9 | O13650 | TFC8/YPL007C | Q12308 |
| SPBC83.10 | emc7 | O94694 | SOP4/YJL192C | P39543 |
| SPBC16C6.01c | | O42925 | RKM3/YBR03W | P38222 |
| SPAC31G5.09c | spk1 | P27638 | KSS1/YGR040W | P14681 |
| SPBC16E9.09c | erp5 | O14324 | ERP1/YAR002C-A | Q05359 |
| SPBC215.05 | gpd1 | P21696 | GPD2/YOL059W | P41911 |
| SPBC609.01 | | O94525 | SSD1/YDR293C | P24276 |
| SPCC757.09c | rnc1 | O74919 | YBL032W/HEK2 | P38199 |
| SPAC24B11.12c | dnf2 | Q09891 | DNF1/YER166W | P32660 |
| SPBC16A3.17c | | O42922 | VBA5/YKR105C | P36172 |
| SPBC27B12.12c | | O13657 | ALR2/YFL050C | P43553 |
| SPBC29A10.11c | vps902 | O94388 | MUK1/YPL070W | Q02866 |
| SPAC19D5.02c | pex22 | Q1K9B6 | EMC10/YDR056C | Q12025 |
| SPCC290.04 | ams2 | Q9URT4 | YMR179W/SPT21 | P35209 |
